# Gas-phase environment activates an alternative catabolic route in toluene-degrading *Acinetobacter*

**DOI:** 10.64898/2026.03.27.714732

**Authors:** Shori Inoue, Shogo Yoshimoto, Maiko Hattori, Hanayo Nakanishi, Yuki Ohara, Katsutoshi Hori

**Affiliations:** Department of Biomolecular Engineering, Graduate School of Engineering, Nagoya University, Furo-cho, Chikusa-ku, Nagoya, Aichi 464-8603, Japan; Friend Microbe Inc., Chikusa-ku, Nagoya, Aichi 464-0858, Japan

## Abstract

Volatile aromatic compounds are important industrial feedstocks but also major environmental pollutants, highlighting the need for bioprocesses for their removal and valorization. Although gas-phase bioprocesses offer practical advantages for handling poorly water-soluble and highly volatile substrates, how gas-phase environments alter microbial metabolism remains poorly understood. Here, we investigated the effect of gas-phase conditions on toluene metabolism in the highly adhesive aromatic hydrocarbon-degrading bacterium *Acinetobacter* sp. Tol 5. A mutant lacking *todC1*, which encodes an essential component of the toluene dioxygenase, failed to grow on toluene in liquid culture but retained the ability to grow on solid media under a toluene atmosphere. Consistent with this phenotype, the mutant showed no detectable toluene degradation in the liquid phase, whereas it degraded toluene under gas-phase conditions after a prolonged lag phase. Gas chromatography–mass spectrometry (GC–MS) analysis revealed the accumulation of *o*-cresol and *p*-cresol specifically in the mutant under toluene vapor, indicating that toluene metabolism had shifted to an alternative route involving cresol intermediates. In addition, transcriptome analysis identified strong induction of the *mph* operon encoding phenol monooxygenase (PMO), suggesting that PMO is a likely candidate enzyme mediating TDO-independent toluene oxidation under gas-phase conditions. Together, these results demonstrate that the gas-phase environment can activate an alternative catabolic route in Tol 5 that is not active during conventional liquid cultivation. Our findings highlight the importance of direct metabolic analysis under gas-phase conditions for understanding and designing bioprocesses using highly volatile substrates.

## Introduction

Aromatic compounds are essential to modern society, serving as industrial solvents, chemical feedstocks, and precursors for pharmaceuticals and polymeric materials. At the same time, they are also major constituents of industrial wastes and environmental pollutants, creating a strong demand for technologies that enable both their removal and productive reuse. Microbial metabolism has long provided a basis for such efforts, and the catabolic capacity of microorganisms has been exploited for the treatment of aromatic pollutants, including volatile compounds, through biofiltration and related processes (1, 2). More recently, advances in metabolic engineering have expanded this concept beyond detoxification, allowing aromatic compounds from wastes or renewable resources to be converted into fuels and value-added chemicals.

Aerobic metabolism of aromatic hydrocarbons begins with activation of the chemically stable aromatic ring. This initial step is typically mediated by multicomponent oxygenases that introduce hydroxyl groups with high chemo- and regioselectivity, thereby producing intermediates that can be funneled into downstream catabolic pathways (3, 4). In many bacteria, the genes required for these initial oxidation steps are organized as a single operon or tightly linked gene cluster and are regulated by dedicated transcriptional regulators. Such genetic and regulatory organization enables efficient substrate-responsive induction while minimizing unnecessary enzyme production and preventing the accumulation of toxic or dead-end intermediates (5–7). This modular architecture has also made aromatic catabolic systems attractive platforms for metabolic engineering in certain soil bacteria such as *Pseudomonas putida* and *Acinetobacter baylyi* ADP1 (8–11). In particular, an engineered alkylbenzene-tolerant *Pseudomonas* strain has enabled the conversion of aromatic hydrocarbons such as benzene and toluene into *cis*,*cis*-muconate (12). These examples demonstrate the significant potential of utilizing microorganisms for the bioconversion of aromatic compounds. However, the efficiency of conventional liquid-phase bioprocesses utilizing these aromatic compounds is frequently hampered by the poor mass transfer of oxygen and volatile, poorly water-soluble aromatic substrates into the culture medium (2, 13, 14).

*Acinetobacter* sp. Tol 5 has emerged as a promising chassis for bioprocesses utilizing such aromatic compounds. Tol 5 possesses a broad aromatic catabolic capacity specifically characterized by the presence of toluene dioxygenase (TDO), which is a rare trait in *Acinetobacter* (15, 16). In addition to its metabolic capability, through the cell-surface nanofiber protein AtaA, Tol 5 exhibits both strong autoaggregation and remarkable adhesiveness, enabling the rapid and robust immobilization of large numbers of cells on solid surfaces without the need for gels or chemical crosslinkers (17, 18). Utilizing this unique adhesiveness, Tol 5 can be applied to gas-phase bioprocesses that enable the efficient supply of oxygen and poorly water-soluble volatile substrates in the absence of bulk water. Indeed, this approach has successfully facilitated the efficient bioconversion of substrates such as toluene and monoterpenes (18, 19). Recent studies have revealed that Tol 5 also demonstrates exceptional tolerance to desiccation (20), making it a robust chassis cell for gas-phase bioproduction. Despite these advantageous traits of Tol 5 for gas-phase reactions, its aromatic metabolism under gas-phase conditions remains largely unexplored.

In this study, we examined toluene metabolism in *Acinetobacter* sp. Tol 5 to determine how liquid-phase and gas-phase conditions influence the metabolic pathway. Using a mutant defective in the previously identified TDO-dependent pathway, we compared toluene utilization under liquid-phase and gas-phase conditions. In addition, GC–MS-based metabolite profiling and transcriptome analysis were performed to characterize changes in the toluene metabolic pathway under gas-phase conditions.

## Materials and methods

### Bacterial strains and culture conditions

The strains used in this study are shown in Table S1. *Escherichia coli* and its transformants were cultured in Luria–Bertani (LB) medium (20066-95; Nacalai Tesque, Kyoto, Japan) at 37 °C. Tol 5 and its mutants were cultured in LB medium or basal salt (BS) medium (21) at 28 °C.

### Gene deletion

The *todC1* deletion mutant was generated by conventional homologous recombination according to a previously reported method (22). The *todF-H* deletion mutant was constructed using a modified procedure based on a combination of Cas9- and recombinase-mediated genome editing (23) and FLP-FRT recombination (24). Gene deletions were verified by cyclodextrin-assisted colony PCR (25) using the primer sets listed in Table S2, followed by DNA sequencing of the amplified target loci.

### Bacterial cell growth

The Tol 5 mutant strains were cultured in 5 mL of LB medium supplemented with 1.2 µL of toluene, *o*-cresol, *m*-cresol, or *p*-cresol at 28 °C for 18 h. The cells were harvested, washed three times with BS medium, and resuspended in BS medium to an optical density at 660 nm (OD_660_) of 1.0.

For liquid culture on toluene, a 0.25-mL aliquot of the suspension was inoculated into 5 mL of BS medium in a glass test tube. Then, 1.2 μL of toluene was added into the tubes. The test tubes were sealed with Viton rubber stoppers and incubated at 28 °C with shaking at 200 rpm. The optical density (OD) was monitored every 60 min using OD-monitorC&T system (Taitec, Saitama, Japan) during the culture. Statistical significance was assessed using Welch’s *t*-test.

For agar culture with toluene, the cell suspension was streaked onto BS agar plates. The plates were placed in AnaeroPack pouches (A-65; Mitsubishi Gas Chemical), along with tissue paper soaked with 5 µL of toluene and incubated at 28 °C for 1 to 5 days.

For agar culture with cresol, the cell suspension was streaked onto BS agar plates containing 0.024% (v/v) *o*-, *m*-, or *p*-cresol. The plates were sealed in the AnaeroPack pouches and incubated at 28 °C for 1 to 5 days.

### Toluene degradation assay

The Tol 5 mutant strains, pre-cultured in 5 mL of LB medium supplemented with 1.2 µL of toluene, were inoculated into 100 mL of LB medium containing 25 µL of toluene and incubated for 18 h. For the liquid-phase reaction, the culture was washed with BS-N medium and resuspended in BS-N medium (21) to an OD_660_ of 1.0. For the gas-phase reaction, an aliquot of the culture equivalent to 10 mL of a cell suspension at an OD_660_ of 1.0 was collected and immobilized onto a glass filter via vacuum filtration. The immobilized cells were washed twice with phosphate-buffered saline (PBS) to remove residual medium components. Either a 10-mL aliquot of the prepared cell suspension or the glass filter containing the immobilized cells was placed into a 50-mL glass vial. The vials were tightly sealed with Teflon-lined stoppers, and 5 µL of toluene was injected into each vial using a microsyringe. The vials for the liquid-phase reaction were incubated at 28 °C with shaking at 115 rpm, whereas those for the gas-phase reaction were incubated statically at 28 °C.

Toluene concentration was monitored using a gas chromatograph equipped with a flame ionization detector (GC-FID; GC-17A, Shimadzu). Chromatographic separation was achieved using an Rtx-200 GC Capillary Column (30 m length × 0.32 mm inner diameter; Restek, Pennsylvania, USA) with nitrogen as the carrier gas. A 50-µL aliquot of the headspace gas was sampled from the vial using a gas-tight syringe (MS-GF100; Ito Corporation, Shizuoka, Japan) and injected into the GC-FID system. The column oven temperature was initially held at 40 °C for 1 min, and then increased to 80 °C at a rate of 20 °C/min. The split ratio was set to 5:1, and the carrier gas flow rate was maintained at 47 mL/min.

### Extraction of metabolites from agar plates

The BS plates inoculated with the Tol 5 mutant strains were placed in tightly sealed AnaeroPack pouches along with tissue paper soaked with 5 µL of toluene, and incubated at 28 °C for 2 days. Following incubation, 5 mL of BS medium was dropped onto each plate, and the surface was gently scraped with a sterile glass spreader to remove the bacterial cells. This washing step was repeated three times. The remaining agar medium was fragmented using a spatula and lyophilized for 2 days. To extract the intermediate metabolites, 15 mL of methanol containing 3 mg/L hexadecane (as an internal standard) was added to the lyophilized agar. The mixture was acidified to approximately pH 2.0 with formic acid, vortexed for 1 min, and centrifuged at 15,000 × g for 5 min. A 1-mL aliquot of the resulting supernatant was filtered for subsequent analysis.

### Gas chromatography–mass spectrometry (GC–MS) analysis

A 1-µL aliquot of each sample was analyzed using a GC-2030 gas chromatograph coupled with a GCMS-QP2020 NX mass spectrometer (Shimadzu, Kyoto, Japan). Chromatographic separation was achieved on an Rtx-200 GC Capillary Column (30 m length × 0.32 mm inner diameter, 0.50 μm film thickness; Restek). Helium was used as the carrier gas. The column oven temperature was initially held at 80 °C for 3 min, increased to 250 °C at a rate of 20 °C/min, and then held at 250 °C for 2 min. The temperatures of the injection port, interface, and ion source were set to 200 °C, 250 °C, and 250 °C, respectively. Mass spectrometry was conducted using electron ionization (EI) at an ionization voltage of 70 eV. Measurements were performed in scan mode with a mass range of *m/z* 35-500. Data acquisition and analysis were performed using LabSolutions GCMS software (Shimadzu).

### RNA sequencing

The Δ*todC* mutant strain was streaked onto BS agar plates supplemented with 0.2% lactic acid and incubated at 28 °C. In parallel, BS plates inoculated with the Tol 5 mutant strains were placed in tightly sealed AnaeroPack pouches along with tissue paper soaked with 5 µL of toluene, and incubated at 28 °C. On the third day, an additional 5 µL of toluene was supplied, and the incubation was continued for a total of 5 days. Subsequently, the cells were harvested by washing the agar surface three times with BS medium, followed by centrifugation (3,000 × g, 25 °C, 10 min). For each growth condition, three biological replicates were prepared. Total RNA was extracted using the Cica Geneus RNA Prep Kit (for Tissue) 2 (Kanto Chemical, Tokyo, Japan) according to the manufacturer’s protocol. Ribosomal RNA was removed using the NEBNext rRNA Depletion Kit (Bacteria) (New England Biolabs, Ipswich, MA, USA), and cDNA libraries were generated with the NEBNext Ultra II RNA Library Prep Kit for Illumina (New England Biolabs, Ipswich, MA, USA). The cDNA libraries were sequenced on the Illumina NextSeq550. Raw FASTQ reads were quality-filtered using fastp (version 0.23.2) and mapped to the Tol 5 genome (accession: AP024708 and AP024709) using Bowtie2 (version 2.5.1) with default parameters. Mapped reads were counted using featureCounts (version 2.0.6).

### Differential gene-expression analysis

Raw read counts of CDSs were analyzed in R using the edgeR package (version 3.42.4). Genes with extremely low expression were removed using the filterByExpr function. Normalization was performed using the trimmed mean of M-values (TMM) method implemented in the calcNormFactors function. Differential gene-expression analysis was conducted using the quasi-likelihood pipeline. Dispersion estimates were obtained using the estimateDisp function, and model fitting was performed with the glmQLFit function. Differential gene-expression between each condition was assessed using the glmQLFTest function.

## Results

### The TDO-deficient mutant of Tol 5 fails to grow in liquid culture but grows on solid media under a toluene atmosphere

We previously identified the TDO gene cluster as the major toluene metabolic pathway in Tol 5 (16). In that study, a gene-disrupted strain with an introduced stop codon in *todC1*, which encodes a key component of TDO, exhibited a near-complete loss of growth in liquid culture with toluene as the sole carbon source. Unexpectedly, however, this strain was still able to grow on agar plates under a toluene atmosphere, suggesting that Tol 5 may possess an alternative gene set or pathway involved in toluene metabolism that can compensate for the loss of TDO. However, sequence analysis of cells collected from these colonies revealed that the population contained revertants in which the stop-codon mutation had been restored. As a result, the toluene assimilation phenotype of the mutant could not be evaluated accurately. Therefore, in the present study, we newly constructed a complete deletion mutant of *todC1* (Δ*todC1*) using the Tol 5_REK_ strain (26). Tol 5_REK_ carries mutations in the restriction-modification system, making genetic manipulation easier, and harbors a mutation in AtaA, resulting in decreased cell adhesion and autoaggregation, thereby improving the accuracy of growth evaluation through turbidity measurements.

When Δ*todC1* was cultivated in liquid medium with toluene as the sole carbon source, no growth was observed even after 150 h of incubation (Fig. 1A). This result, consistent with our previous findings, indicates that *todC1* is essential for growth on toluene in liquid culture. In contrast, the deletion mutant clearly grew on BS agar plates placed under a toluene atmosphere, whereas no colonies were formed on the same medium in the absence of toluene (Fig. 1B). This result indicates that Δ*todC1* is able to utilize toluene as a carbon source on solid media. These results suggest that, although TDO is the major pathway for toluene metabolism in liquid culture, Tol 5 possesses an alternative gene or pathway that can compensate for the loss of TDO during growth on solid media under a toluene atmosphere.

**Figure 1.**
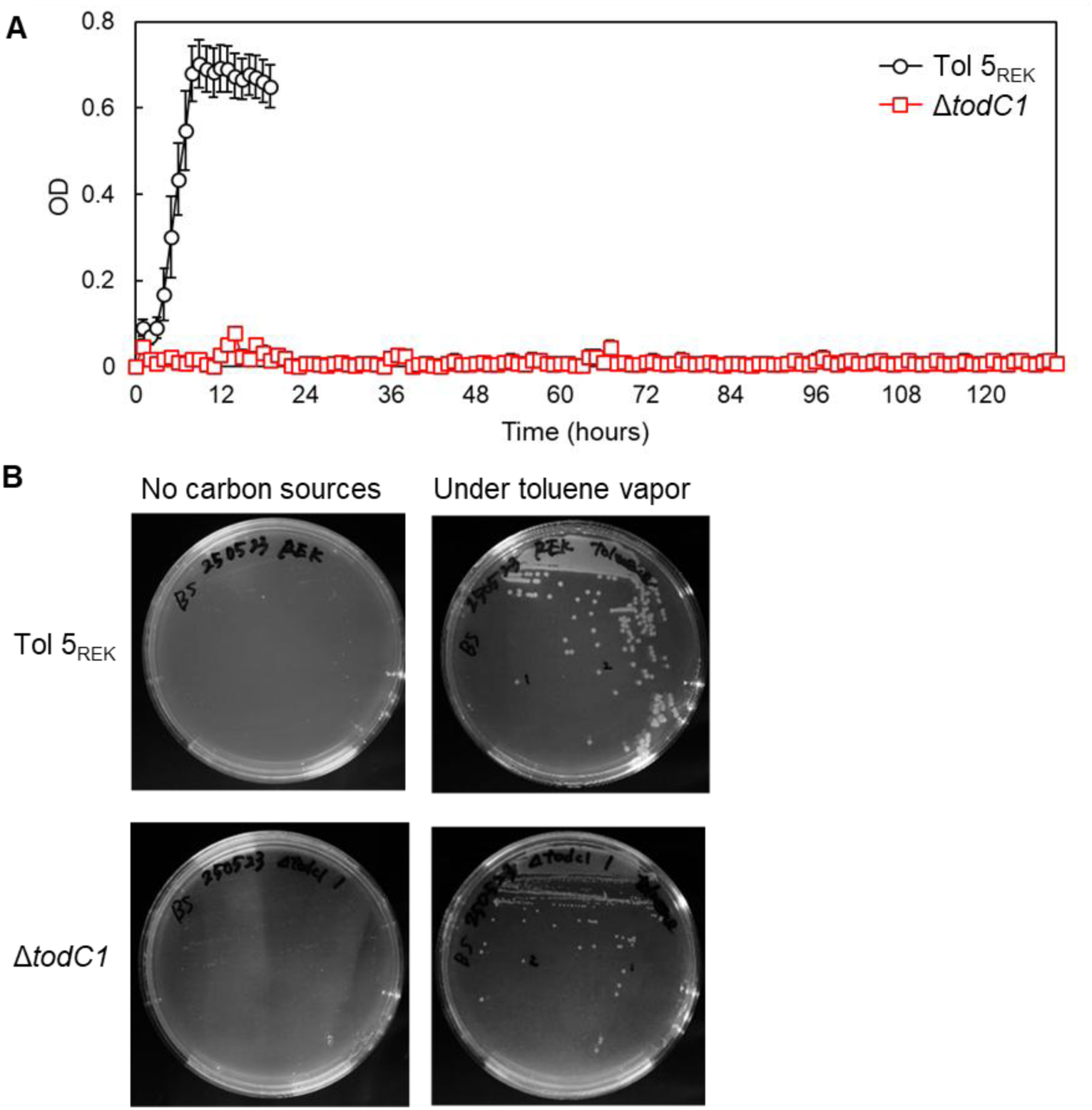
Growth of Tol 5_REK_ and Δ*todC1* mutant strain. (A) Tol 5_REK_ and the Δ*todC1* mutant strain were inoculated to BS medium containing toluene as a sole carbon source. The data are presented as mean ± SD (n = 3). (B) Tol 5_REK_ and the Δ*todC1* mutant were inoculated onto BS agar plates and incubated either in air (left) or under a toluene vapor atmosphere (right).

### Gas-phase environment is a key factor enabling toluene metabolism in the TDO-deficient mutant

In liquid culture, toluene is supplied as a dissolved substrate in the medium, whereas on agar plates it is supplied in the gaseous state as vapor. We previously reported that, under such gas-phase conditions, microorganisms exhibit metabolic profiles and gene expression patterns distinct from those observed in liquid-phase environments (27). Based on this, we hypothesized that the difference in toluene metabolism observed between liquid and solid cultivation might be attributable, at least in part, to the effect of the gas-phase environment. To test this hypothesis, Tol 5_REK_ and Δ*todC1* cells grown in LB medium were subjected to toluene degradation assays either in nitrogen-free BS medium (BS-N) under liquid-phase conditions or under gas-phase conditions. In the gas-phase assay, cells were immobilized on glass filters, and no agar or liquid medium was present.

Under liquid-phase conditions, Tol 5_REK_ degraded toluene within 12 h, whereas Δ*todC1* showed no detectable toluene degradation even after 150 h (Fig. 2A). This result again indicates that TDO is essential for toluene degradation under liquid-phase conditions. In contrast, under gas-phase conditions, Δ*todC1* degraded toluene after a prolonged lag phase of approximately 100 h (Fig. 2B). This result indicates that Δ*todC1* can activate a toluene degradation pathway that compensates for the loss of TDO only under gas-phase conditions, identifying the gas-phase environment as a key factor governing the change in toluene metabolism.

**Figure 2.**
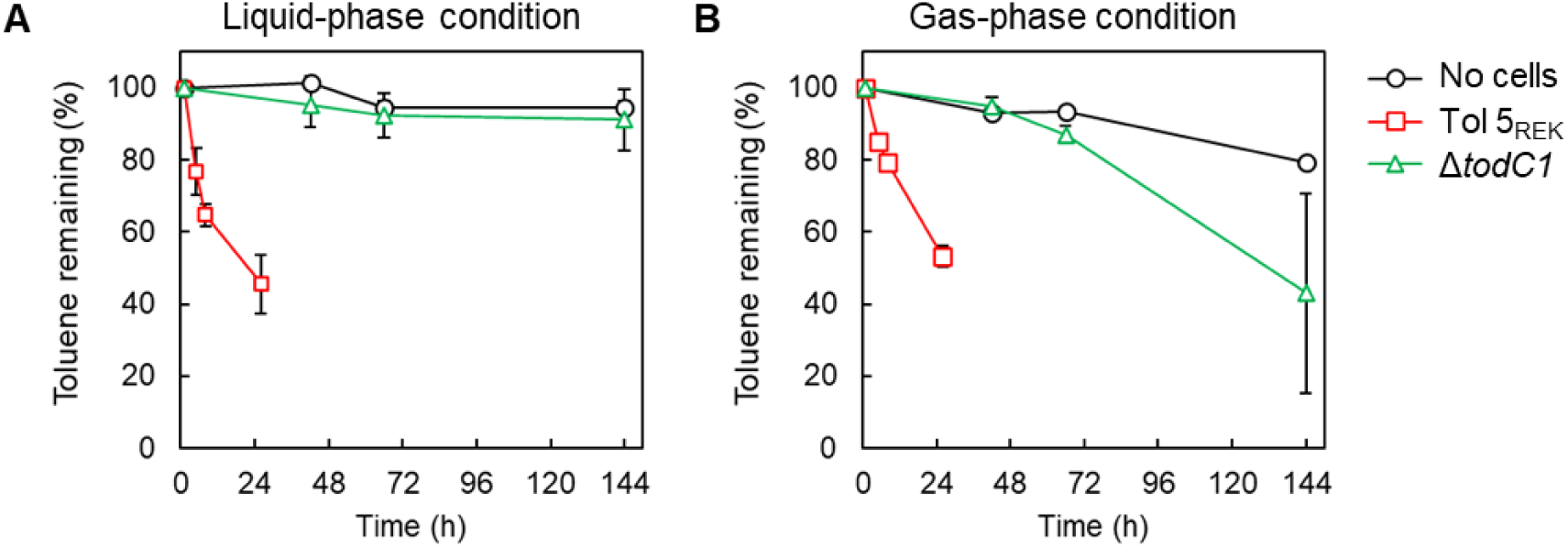
Toluene degradation by the Tol 5_REK_ and Δ*todC1* mutant strain. Cells were incubated with toluene in (A) the liquid phase (BS-N medium) or (B) the gas phase. Toluene concentration in the headspace was quantified using GC-FID. The vertical axis represents the percentage of remaining toluene relative to the amount at 1 hour after toluene addition. The data are presented as mean ± SD (n = 3).

### Toluene degradation by Δ*todC1* is accompanied by cresol accumulation

To clarify the alternative toluene metabolic pathway responsible for toluene metabolism under gas-phase conditions, metabolites were extracted from carriers or agar media after exposure of Tol 5_REK_ and Δ*todC1* to toluene vapor, and analyzed by GC–MS. Comparison of the GC–MS chromatograms revealed two peaks that were detected only in Δ*todC1* in the presence of toluene (Fig. 3A), suggesting that these compounds were toluene metabolites. Analysis of their mass spectra identified them as *o*-cresol and *p*-cresol, respectively (Fig. 3B). Because these metabolites were not detected in the non-mutant Tol 5_REK_ strain, cresols are likely to be specifically associated with the alternative toluene metabolism operating in Δ*todC1* under gas-phase conditions.

**Figure 3.**
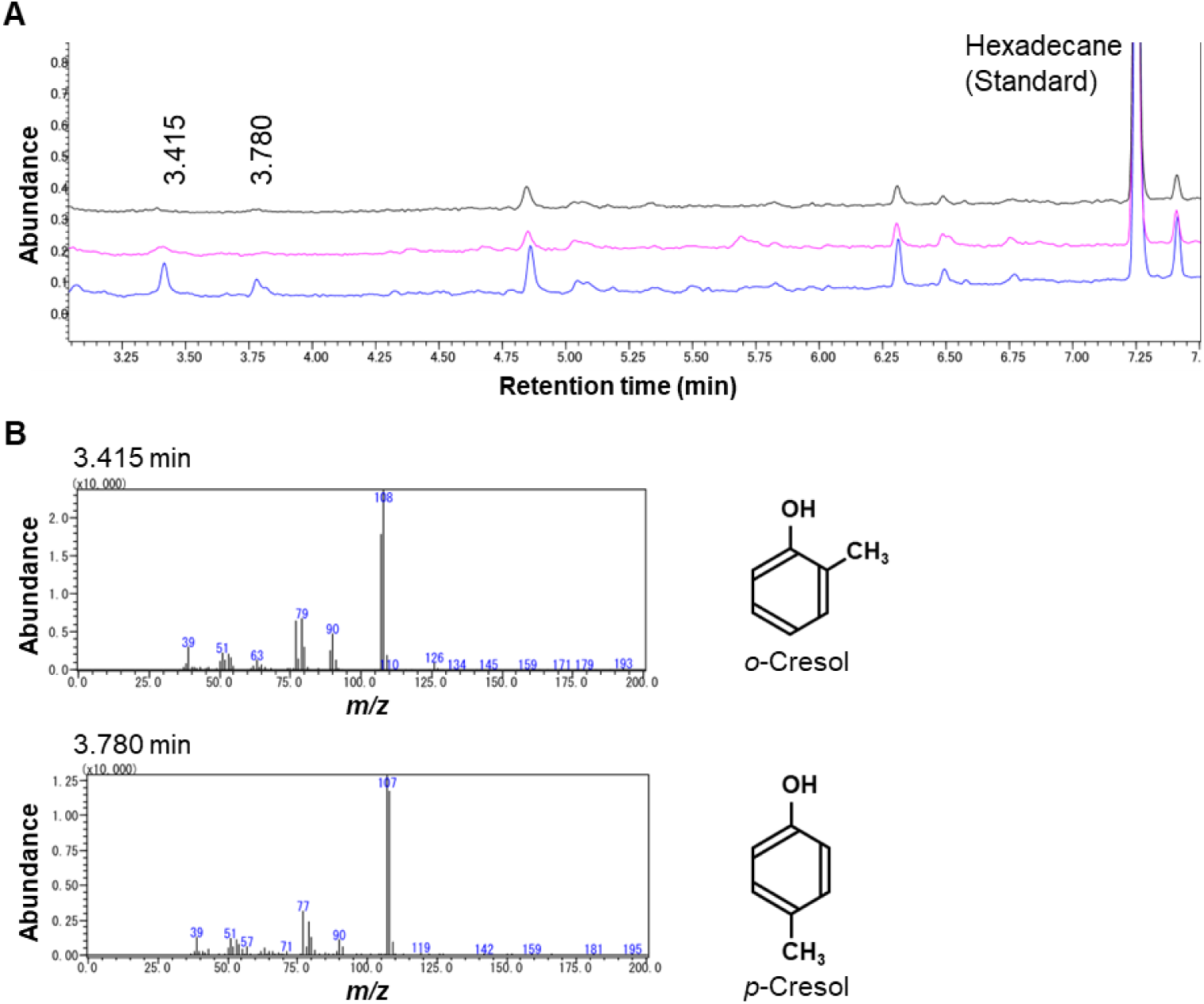
GC–MS analysis of metabolites accumulated during growth on agar plates under toluene vapor. (A) Chromatograms of metabolites extracted from the agar plates. The chromatograms represent samples extracted from BS plates without any cells (black), with the Tol 5_REK_ (pink), and with the Δ*todC1* mutant (blue). Peaks corresponding to retention times of 3.415 and 3.780 min were detected only in the extract from the Δ*todC1* mutant. Hexadecane was added as an internal standard. (B) Mass spectra of the peaks at 3.415 min and 3.780 min in the chromatogram of the Δ*todC1* mutant. Based on the fragmentation patterns (*m/z*), the metabolites were identified as *o*-cresol and *p*-cresol, respectively.

In the major toluene metabolic pathway of Tol 5, toluene is oxidized by TDO and converted to 3-methylcatechol via a dihydroxylated intermediate, and subsequently cleaved by an oxidative meta-cleavage reaction, and funneled into downstream catabolic pathways (Fig. 4A). The detection of cresols in gas-phase Δ*todC1* indicates that, rather than being restored through complementation of TodC1 function, toluene metabolism is redirected to a distinct pathway in which cresols serve as intermediates. Since Δ*todC1* was able to grow on agar plates under a toluene atmosphere, these cresols are likely further metabolized and funneled into central metabolism. To test whether cresol isomers can indeed serve as sole carbon sources, Tol 5_REK_ and Δ*todC1* were cultivated on agar plates containing individual cresol isomers. Both strains grew on *o*-cresol and *m*-cresol, but not on *p*-cresol (Fig. 4B). Given that hydroxylation of both *o*-cresol and *m*-cresol by aromatic hydroxylases is known to yield 3-methylcatechol (28, 29), these findings suggest that both cresol isomers are metabolized via 3-methylcatechol before entering downstream metabolism. Consistent with this, in Tol 5, a strain lacking these downstream *tod* genes (Δ*todF-H*) failed to grow on any of the cresol isomers (Fig. 4B). These results indicate that the *tod* genes are essential for cresol metabolism in Tol 5. Based on these findings, we propose the following toluene metabolic route for gas-phase Δ*todC1* (Fig. 4A). First, toluene is converted to *o*- and *m*-cresol by an as-yet-unidentified aromatic monooxygenase that functions specifically under gas-phase conditions. These cresols are then further hydroxylated to 3-methylcatechol, which is subsequently converted to pyruvate and acetate by the native Tod enzymes.

**Figure 4.**
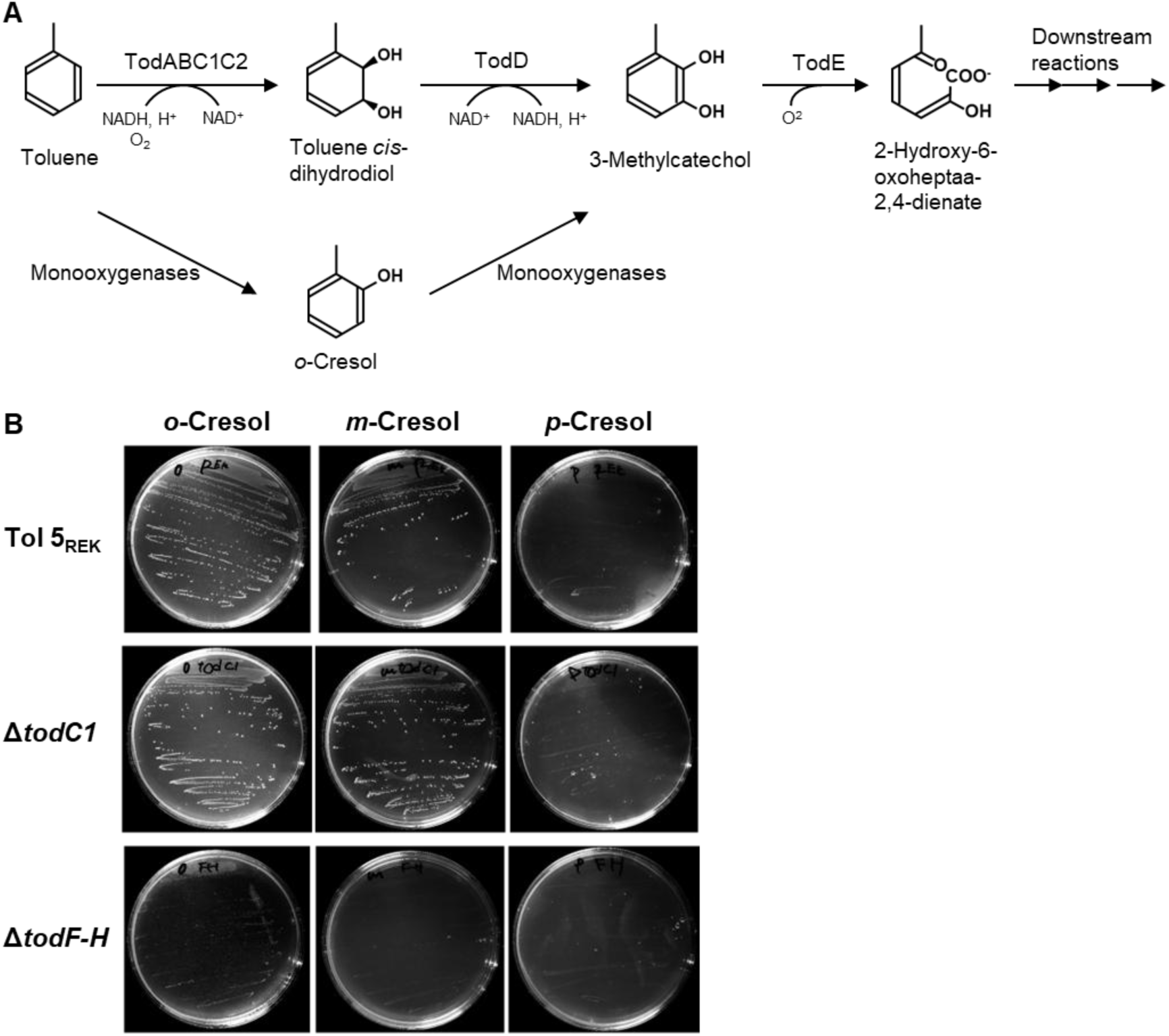
Proposed metabolic pathway for toluene degradation and cresol assimilation by Tol 5 mutant strains. (A) Schematic representation of the proposed toluene degradation pathways. The upper route indicates the canonical TDO-dependent pathway, whereas the lower route indicates the newly proposed alternative pathway via *o*-cresol mediated by unidentified monooxygenases. (B) Tol 5_REK_, Δ*todC1*, and Δ*todF-H* mutants were inoculated onto BS agar plates supplemented with *o*-, *m*-, or *p*-cresol.

### Transcriptomic identification of candidate genes for TDO-independent toluene metabolism

To identify genes involved in the alternative pathway that compensates for the loss of TDO during toluene metabolism in Δ*todC1*, we performed transcriptome analysis. Gene expression in Δ*todC1* cells grown on agar plates with toluene vapor was compared with that in cells grown with lactate as the sole carbon source (Fig. 5A). Numerous differentially expressed genes were detected (|logFC| > 1, logCPM > 1, FDR < 0.01), including several that were both highly abundant and strongly upregulated under the toluene atmosphere. Among the 18 most strongly induced genes (logFC > 7), we identified not only the *tod* operon, which encodes the major toluene metabolic pathway in Tol 5, but also genes encoding the components of the phenol monooxygenase (PMO) complex and catechol 1,2-dioxygenase (Table 1). Because the *mph* genes encoding PMO and *catA*, encoding catechol 1,2-dioxygenase, are located in the same operon (15), these data indicate strong induction of the *mph* operon. These findings suggest that PMO is a likely candidate for the enzyme that compensates for TDO in toluene conversion.

**Figure 5.**
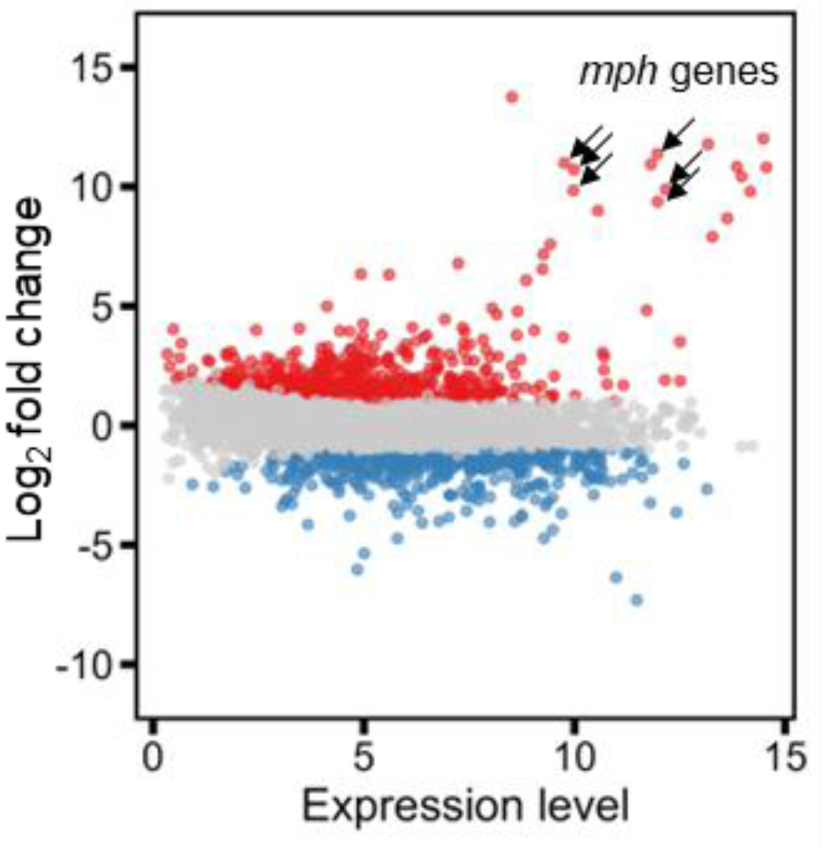
MA plot of gene expression changes in the Δ*todC1* mutant. MA plot comparing gene expression in the Δ*todC1* mutant cultured on BS agar plates under toluene vapor relative to growth on sodium DL-lactate. The vertical axis represents the logFC (toluene / lactate), and the horizontal axis represents the logCPM. Red and blue dots denote significantly up-regulated (logFC > 1, logCPM > 1, FDR < 0.01) and down-regulated genes (logFC < −1, logCPM > 1, FDR < 0.01), respectively. Gray dots indicate genes with no significant differential expression. Arrows point to the *mph* genes encoding the components of PMO.

**Table 1.**
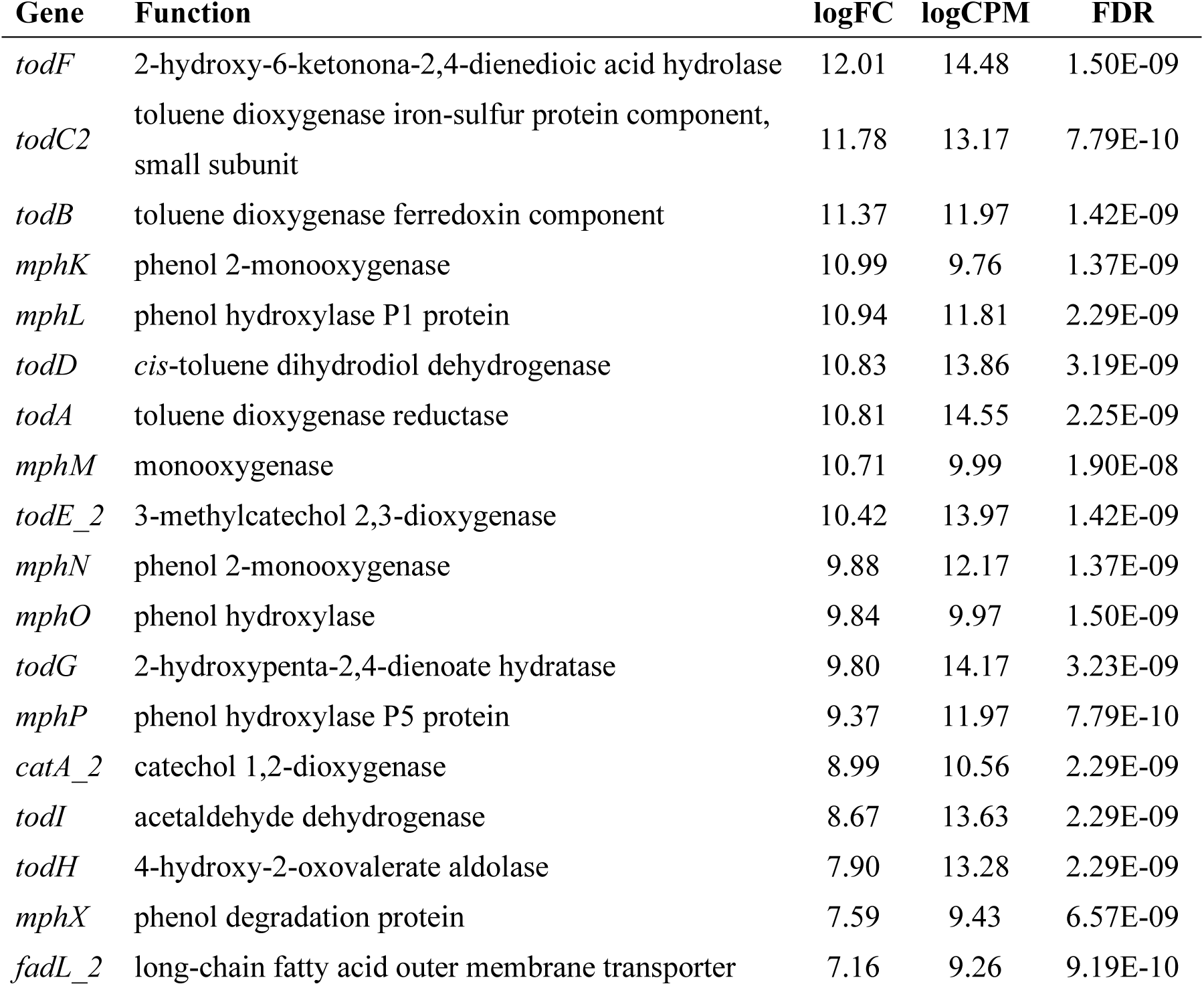
Top 18 highly upregulated genes in the Δ*todC1* mutant grown on toluene.

| Gene | Function | logFC | logCPM | FDR |
| --- | --- | --- | --- | --- |
| *todF* | 2-hydroxy-6-ketono-2,4-dienedioic acid hydrolase | 12.01 | 14.48 | 1.50E-09 |
| *todC2* | toluene dioxygenase iron-sulfur protein component, small subunit | 11.78 | 13.17 | 7.79E-10 |
| *todB* | toluene dioxygenase ferredoxin component | 11.37 | 11.97 | 1.42E-09 |
| *mphK* | phenol 2-monooxygenase | 10.99 | 9.76 | 1.37E-09 |
| *mphL* | phenol hydroxylase P1 protein | 10.94 | 11.81 | 2.29E-09 |
| *todD* | *cis*-toluene dihydrodiol dehydrogenase | 10.83 | 13.86 | 3.19E-09 |
| *todA* | toluene dioxygenase reductase | 10.81 | 14.55 | 2.25E-09 |
| *mphM* | monooxygenase | 10.71 | 9.99 | 1.90E-08 |
| *todE_2* | 3-methylcatechol 2,3-dioxygenase | 10.42 | 13.97 | 1.42E-09 |
| *mphN* | phenol 2-monooxygenase | 9.88 | 12.17 | 1.37E-09 |
| *mphO* | phenol hydroxylase | 9.84 | 9.97 | 1.50E-09 |
| *todG* | 2-hydroxypenta-2,4-dienoate hydratase | 9.80 | 14.17 | 3.23E-09 |
| *mphP* | phenol hydroxylase P5 protein | 9.37 | 11.97 | 7.79E-10 |
| *catA_2* | catechol 1,2-dioxygenase | 8.99 | 10.56 | 2.29E-09 |
| *todI* | acetaldehyde dehydrogenase | 8.67 | 13.63 | 2.29E-09 |
| *todH* | 4-hydroxy-2-oxovalerate aldolase | 7.90 | 13.28 | 2.29E-09 |
| *mphX* | phenol degradation protein | 7.59 | 9.43 | 6.57E-09 |
| *fadL_2* | long-chain fatty acid outer membrane transporter | 7.16 | 9.26 | 9.19E-10 |

## Discussion

Aromatic compounds are indispensable to a wide range of industries, but they are also major industrial wastes and environmental pollutants. In particular, alkylbenzenes and related aromatic compounds are chemically highly stable because of the large resonance energy of their aromatic rings and are therefore generally recalcitrant in natural environments. Some microorganisms, however, can exploit powerful and complex oxidative enzymes to degrade these compounds and convert them into simpler biomolecules for growth and metabolism (4, 30). In this study, we found that a Tol 5 mutant lacking the TDO gene remained capable of growth on agar plates and demonstrated that an alternative toluene catabolic pathway is activated specifically under gas-phase conditions. Although several studies have highlighted the advantages of gas-phase reactions for bioprocess applications (19, 31, 32), to our knowledge, this study provides the first example showing that a gas-phase-induced metabolic shift can be essential for carbon source utilization. This finding indicates that the design of gas-phase bioprocesses requires not only insights gained from conventional liquid-culture studies but also direct metabolic analyses under gas-phase conditions. In addition, aromatic oxidation enzymes are known to catalyze regioselective hydroxylation of a broad range of allyl compounds, and their application in bioprocesses for the selective oxidation of monoterpenes into value-added products has attracted increasing attention (33). Because many of these substrates are highly volatile, gas-phase reactions have also been actively investigated for such conversions (31). Our findings further suggest that gas-phase-specific shifts in metabolic conversion should be taken into account when designing gas-phase bioprocesses for highly volatile compounds, including but not limited to aromatic substrates.

We previously reported that the major toluene catabolic pathway in Tol 5 is mediated by TDO encoded by the *tod* operon (16). In this pathway, toluene is first converted to a dihydrodiol by TDO and then transformed into 3-methylcatechol through the action of a dehydrogenase. In the present study, however, cresol accumulation was detected in the TDO-deficient strain metabolizing toluene under the gas-phase condition (Fig. 3A and B). Furthermore, differential gene expression analysis identified PMO as a candidate monooxygenase responsible for a two-step hydroxylation pathway in which cresol serves as an intermediate (Table 1). PMO has been reported to hydroxylate not only phenol but also *o*-, *m*-, and *p*-cresol, as well as benzene (29). In addition, *E. coli* expressing the PMO gene from *Arthrobacter* sp. W1 was shown to catalyze both oxidation steps from toluene to cresol and from cresol to 3-methylcatechol (28). Taken together, these findings suggest that PMO likely bypasses the canonical TDO-dependent pathway in Tol 5 under gas-phase conditions by catalyzing these two sequential reactions. A key difference between toluene conversion via the TDO and via the PMO lies in their requirements for reducing power and oxygen. In the TDO-dependent pathway, molecular oxygen is activated by reducing equivalents derived from NADH to convert the aromatic ring of toluene into a diol, whereas an equimolar amount of NADH is regenerated in the subsequent dehydrogenation step. In contrast, the two-step hydroxylation pathway mediated by PMO consumes two molecules each of NADH and oxygen. Accordingly, the preferential use of the more energy-efficient TDO-dependent pathway by the wild-type strain during toluene metabolism appears physiologically reasonable, despite the presence of this alternative PMO-mediated route in Tol 5. Under conditions in which TDO is nonfunctional, however, activation of PMO may provide an alternative route for toluene oxidation, thereby contributing to the robustness of aromatic compound metabolism in Tol 5.

The shift in the toluene metabolic pathway observed under gas-phase conditions in this study appears to be associated with the induction of the *mph* genes encoding PMO. In Tol 5, the PMO genes are organized as a single operon and are thought to be transcriptionally regulated by the adjacent regulatory factors *mphR* and *mphX* (15). A previous study reported that, during liquid cultivation with toluene as the carbon source, the *tod* operon encoding TDO and its downstream enzymes was strongly activated, whereas the *mph* operon encoding PMO was only minimally induced (34). In addition, *mphR* is homologous to *mopR* from *Acinetobacter calcoaceticus*, and MopR has been shown in vitro to bind a broad range of hydroxybenzenes but not toluene (34). These observations suggest that induction of PMO under gas-phase conditions is likely governed by a transcriptional regulatory mechanism distinct from that observed during conventional toluene cultivation. Such regulation may be triggered by multiple factors associated with the absence of bulk water, including stress responses, changes in oxygen or substrate availability, and the accumulation of metabolic intermediates. Further molecular investigation will be required to clarify which factors drive this gas-phase-specific transcriptional shift.

In conclusion, this study shows that the gas-phase environment can reshape toluene metabolism in *Acinetobacter* sp. Tol 5 by activating an alternative catabolic route, likely involving PMO. This finding identifies the reaction environment itself as an important determinant of pathway selection in aromatic hydrocarbon metabolism and further indicates that metabolic behavior inferred from conventional liquid cultivation cannot always be directly extrapolated to gas-phase systems. Together, these insights provide a conceptual basis for understanding and designing gas-phase bioprocesses for highly volatile substrates under water-limited conditions.

## Acknowledgements

We thank Eriko Kawamoto for her technical assistance.

This research was supported by the Graduate Program of Transformative Chem-Bio Research at Nagoya University supported by MEXT (WISE Program) to SI, the Japan Science and Technology Agency (JST) SPRING (Grant Number 8 JPMJSP2125) to SI, the Japan Society for the Promotion of Science (JSPS) KAKENHI (Grant Number JP24H00043) to KH and SY.

## Supporting Information

**Table S1.** Bacterial strains used in this study.

| Strain | Description | Reference |
| --- | --- | --- |
| *Acinetobacter* sp. |  |  |
| Tol 5 | Wild type strain | (35) |
| Tol 5<sub>REK</sub> | Restriction enzyme-encoding genes and *ataA* knockout mutant of Tol 5, REK123Δ*ataA* | (26) |
| Tol 5<sub>REK</sub> Δ*todC1* | *TodC1*-deletion mutant of Tol 5<sub>REK</sub> | This study |
| Tol 5<sub>REK</sub> Δ*todF-H* | *todF,C1,C2,B,A,D,E,G,I,H*-deletion mutant of Tol 5<sub>REK</sub> | This study |

**Table S2.**
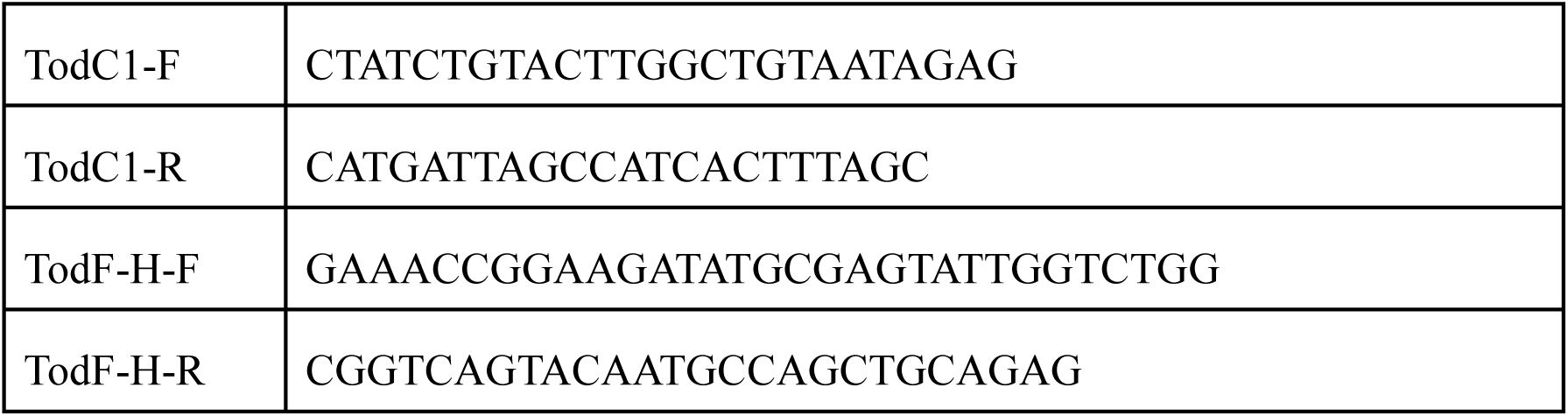
Primers used in this study.

